# Binding of the peptide deformylase on the ribosome surface modulates the structure and dynamics of the exit tunnel interior

**DOI:** 10.1101/2022.04.20.488877

**Authors:** Hugo McGrath, Michaela Černeková, Michal H. Kolář

**Affiliations:** Department of Physical Chemistry, University of Chemistry and Technology, Technická 5, 16628 Prague, Czech Republic

## Abstract

Proteosynthesis on ribosomes is regulated at many levels. Conformational changes of the ribosome, possibly induced by external factors, may transfer over large distances and contribute to the regulation. The molecular principles of this long-distance allostery within the ribosome remain poorly understood. Here, we use structural analysis and atomistic molecular dynamics simulations to investigate peptide deformylase (PDF), an enzyme that binds to the ribosome surface near the ribosomal protein uL22 during translation and chemically modifies the emerging nascent peptide. Our simulations of the entire ribosome–PDF complex reveal that the PDF undergoes a swaying motion on the ribosome surface at the sub-microsecond time scale. We show that the PDF affects the conformational dynamics of parts of the ribosome over distances of more than 5 nm. Using a supervised-learning algorithm we demonstrate that the exit tunnel is influenced by the presence or absence of PDF. Our findings suggest a possible effect of the PDF on the nascent peptide translocation through the ribosome exit tunnel.

## 1 Introduction

During translation, the amino acids are delivered on tRNAs to the peptidyl transferase centre (PTC) deep in the large ribosomal subunit, where they are added one by one to the elongating nascent chain (NC). A 10 nm-long tunnel accommodates the NC before it exits on the ribosome surface into the cytosol. The tunnel is mostly composed of rRNA, but several ribosomal proteins contribute to the tunnel walls as well. In bacteria, the loop extensions of uL4 and uL22 create a tunnel constriction. Closer to the tunnel exit, proteins uL23 and uL24 may affect the co-translational folding of small protein domains.^1^

Altogether, the tunnel walls form an active environment.^2^ Some NC sequences specifically interact with the walls or low-molecular factors in the exit tunnel. Often, these interactions have profound physiological consequences. For instance, translational stalling caused by the SecM peptide affects expression of a downstream gene^3^ and regulates protein transmembrane export. The TnaC sequence interacts with free tryptophan and regulates its catabolism.^4,5^ Erythromycin and other macrolide antibiotics stall the ribosome and through the interactions with certain NCs they regulate the expression of macrolide-resistance genes.^6,7^ The exit tunnel may also induce or facilitate folding of secondary structure motives^8^ or small protein domains.^9^

Several studies have reported signal transfer throughout the tunnel walls. The translational arrest caused by SecM involves an allosteric signaling from the constriction site through the rRNA nucleotide A2062 to the PTC.^10,11^ Likewise, another arresting peptide – VemP – switches off the PTC through a series of NCo o owall and NCo o oNC interactions.^8,12^ In the presence of the ErmBL peptide, the binding of erythromycin was proposed to induce an allosteric change in rRNA nucleotides U2504–U2506 which modulates PTC and contributes to the stalling 6. A number of potential allosteric sites have been recently reviewed in the *Thermus thermophilus* and *Eschericha coli* ribosomes.^13^

The ribosome surface contains binding sites for protein-biogenesis translation factors like the trigger factor, signal recognition particle (SRP) or enzymes ensuring the NC modifications.^14^ A question arises, whether the information about surface status spreads deeper into the ribosome interior, or, vice versa, whether the functional state of the ribosome or NC presence affects the recruitment of translation factors.

Indeed, an allosteric signal has been proposed to modulate SRP-mediated membrane targeting on the bacterial ribosome. The SRP binding affinity to the ribosomal protein uL23 was about 100 times higher, when a short NC was present in the tunnel as compared to a vacant ribosome.^15^ Strikingly, no sequence effect was observed, suggesting a universal conformational change of uL23. A coupling between a loop of uL23 and exit-tunnel nucleotides observed in coarse-grained molecular dynamics (MD) simulations was proposed as a possible mechanism of allosteric signal transfer.^13^

The constriction-site proteins uL4 and uL22 are highly conserved across all three domains of life. uL4 and uL22 were initially postulated as a discrimination gate^16^ but this idea has not been verified. Still, due to a number of charged residues and its vicinity to the PTC, the constriction site likely interacts with many NCs.^17^

The globular part of uL22 is exposed to the ribosome surface and contributes to the binding site of the peptide deformylase (PDF). PDF removes the formyl group from the N-terminal formylmethionine and initiates co-translational maturation of NCs. PDF is essential for eubacteria,^18^ therefore it was proposed as a potential drug target for antibiotics.^19^ A PDF binder actinonin and its analogues were also identified as potential anti-cancer agents^20^ targeting mitochondrial PDF.

The PDF class I present in *E. coli* binds to the ribosome surface through its C-terminal *α*-helix.^21^ The binding interface was described by cryogenic electron microscopy (cryo-EM) at 3.7 A resolution using a model 16-residue peptide from the PDF C-terminus. The full PDF was studied by cryo-EM at lower resolution (10–15 Å)^22^ and it was shown that PDF can interfere with other protein biogenesis factors. The effect of PDF on the ribosome structure and dynamics has not been the center of focus.

Here, we study the interaction of the complete PDF with the ribosome and test the anticipated allostery between the ribosome surface and its interior triggered by the PDF binding. We use all-atom molecular dynamics (MD) simulations, which have proven useful in studying conformational dynamics of the ribosome.^23^ We simulate the entire *E. coli* ribosome in explicit water to compare dynamics of the ribosome in the presence and absence of PDF. We apply a classification supervised-learning algorithm to predict the presence or absence from the conformations of various ribosome parts. Our results suggests that the PDF is highly flexible when bound to the ribosome and that the tunnel walls are modulated by the PDF.

## 2 Results

### 2.1 PDF is highly flexible when bound to the ribosome

We carried out four independent unbiased MD simulations of the ribosome (PDF–) and ribosome-PDF complex (PDF+) and analyzed the last 1 *μ*s of each trajectory. As expected, the PDF remained bound to the ribosome surface throughout the course of all PDF+ simulations. The structural context of PDF is shown in Fig. 1A. The PDF binds through its C-terminal *α*-helix to a groove formed by ribosomal proteins uL22, bL32, and rRNA. The C-terminus of PDF points to the exit of the ribosomal tunnel. The catalytic site of PDF is accessible from the right hand side, as indicated by the black arrow in Fig. 1A.

**Figure 1:**
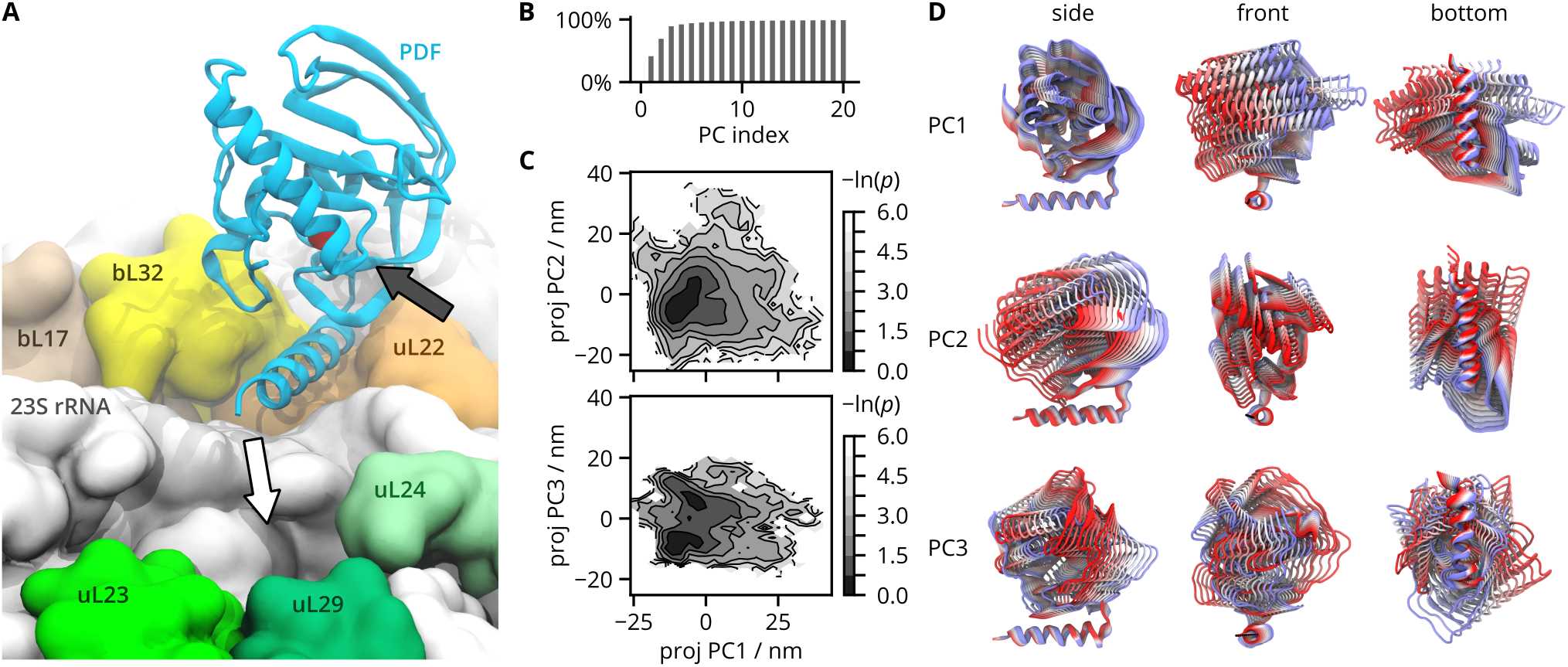
Analysis of the PDF motion on the ribosome surface. A) Structural context of PDF with labeled ribosome components. The white arrow points to the exit of the ribosomal tunnel, the black arrow shows the access to the catalytic site. The catalytic Ni^2+^ ion is shown as a red sphere. B) Percentage of explained total variance as a function of number of PCs. C) Projections of the trajectories onto eigenvectors PC1, PC2, and PC3 are shown as the negative natural logarithm of a probability density *p.* D) Structural representations of PC1, PC2, and PC3. The color scale goes from negative values (red) through white to positive (blue). The extremes represent values observed in MD simulations.

To find the dominant conformational mode of PDF, we used principal component analysis (PCA). PCA was carried out for the PDF backbone atoms after least-square alignment of the large ribosomal subunit C*_α_* and P atoms to a high-resolution cryo-EM structure (PDB id: 5AFI^24^). Therefore, the obtained principal components reflect the PDF motion relative to the large subunit.

From the percentage of explained variance plotted in Fig. 1B, we can see that the first three eigenvectors – principal components (PCs) – cover 90% of the total variance. The conformational modes described by the PCs are shown in Fig. 1D. PC1 is a side-to-side swaying motion of the N-terminal catalytic domain about the C-terminal helix best visible from the front, when looking along the helical axis. The C-terminal helix remains well localized on the ribosome surface and forms the axis of rotation. The last two C-terminal amino acids fluctuate more than the rest of the *α*-helix. PC2 is a front-to-back swaying best visible from a side view. PC3 is a rotational motion about a vertical axis perpendicular to the ribosome surface. Given the trajectory lengths of 1 *μ*s, the swaying of PDF is fast, on the order of hundreds of nanoseconds (time traces of PCs are given in Fig. S1).

Fig. 1C shows 2D projections of the trajectories onto the first three PCs. Projections on PC1 and PC2 show a single broad minimum. The PC3 has two minima which represent left- or right-rotated substates of the C-terminal helix. The motion along PC3 is less extensive than along PC1 or PC2. Overall, the results of PCA display a fast and extensive conformational motion of the PDF on the ribosome.

### 2.2 PDF modulates structure of uL22 inside the ribosomal tunnel

By averaging the trajectory frames over the simulation time, we obtained simulation-derived structural models of PDF+ and PDF– at atomic resolution. For uL22, we calculated residuewise root-mean-square deviation (rwRMSD) between the two models (Fig. 2). The rwRMSD informs us of the effect of PDF on the uL22 structure. Previously, an atom-resolved model of the ribosome with the C-terminal helix of PDF was determined by X-ray crystallography (PDB 4V5B^21^), so we calculated rwRMSD of uL22 between the PDF+ X-ray model and a PDF–cryo-EM model (PDB 5AFI^24^).

**Figure 2:**
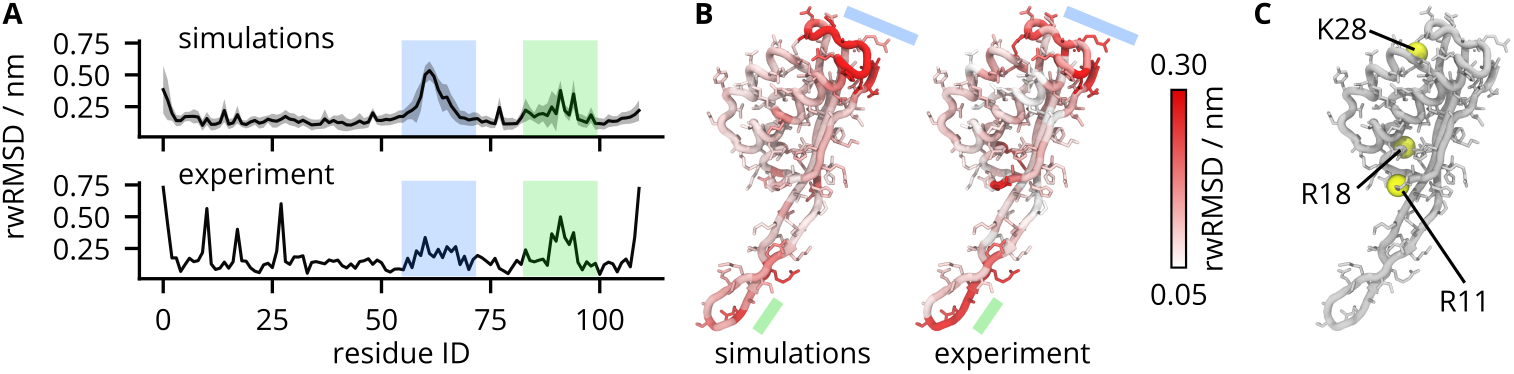
Residue-wise root-mean-square deviation (rwRMSD) of uL22 with and without PDF bound. A) Simulation-based rwRMSD were calculated for average structures from all pairs of PDF+ and PDF– trajectories and averaged (black line). The standard deviations over the pairs are shown as a gray area. Experiment-based rwRMSD represents a single pair of PDF+ (PDB 4V5B^21^) and PDF– (PDB 5AFI^24^). The surface and tunnel loops are highlighted by blue and green, respectively. B) rwRMSD projected onto the uL22 structure without PDF. C) Selected residues with a high experiment-based rwRMSD.

Overall, the simulation and experimental rwRMSDs agree well. Simulation-based rwRMSD reveals two regions of uL22, which are affected by the PDF binding, namely a surface loop between N61 and D67, and the uL22 extension of residues around R92, which protrudes into the exit tunnel (highlighted in blue and green in Figs. 2AB). These regions are also identified by the experiment-based rwRMSD. The experiment-based rwRMSD further suggests that R11 and R18, located in the hinge between the globular part of uL22 and its loop extension, and K28, located on the surface, are sensitive to PDF presence (Fig. 2C). High RMSD of the three residues reflects highly rotated side chains in the PDF+ and PDF– experimental models. Because all simulations were initiated from non-rotated states of R11, R18, K28 in PDF– and the side chain rotations likely require longer time scales, they were not observed in the simulations. All simulations were started from the PDF– conformation of the ribosome. Therefore, the observation that differences between PDF+ and PDF– emerging from the simulations are in agreement with the differences in the experimentally derived structures provide evidence that our simulation time scales are sufficient to capture the differences in dynamics.

### 2.3 Classification Model of the Ribosome Surface Status

From the structural analysis it appears that the tip of uL22 is sensitive to the presence of PDF. To investigate possible allosteric pathways, which could transfer the information between the ribosome surface and its interior, we carried out principle component regression (PCR) of several ribosome parts. PCR is a supervised machine-learning algorithm extending the capabilities of PCA (see Methods).^25^ A binary function *f* labeling the presence (*f* = 1) or absence (*f* = 0) of PDF on the ribosome surface was defined, and a model *f_m_* was trained to predict *f* from a subset of PCs of various parts of the ribosome. Two PDF+ and two PDF– trajectories were used for training and the remaining pairs of trajectories were used for validation. Consequently, each frame from the validation set was classified to belong to PDF+ or PDF– subset based on whether *f_m_* was greater than or lower than 0.5, respectively. We assume that the selections important for a good prediction are those which change due to presence of the PDF.

Indeed, investigating various ribosome parts by PCR showed that some atom selections (their PCs) can predict the presence or absence of PDF better than others (Fig. 3A). Among the ribosomal proteins near the PDF binding site, only uL22 yielded a good model with Pearson correlation coefficient *R* = 0.92 ± 0.04 (standard deviation over validation sets, see Methods). The model constructed from uL22 PCs classified 98% of frames from the validation set correctly. As a control, we selected all 20th residues of ribosomal proteins of the large subunit. They are spread around the whole subunit so their conformations should not correlate with the presence of PDF on the surface. Indeed, poor correlation between *f_m_* and *f* was observed for this selection (R = 0.21 ± 0.12) and only 55% of PDF+ and 64% of PDF– frames were correctly classified.

**Figure 3:**
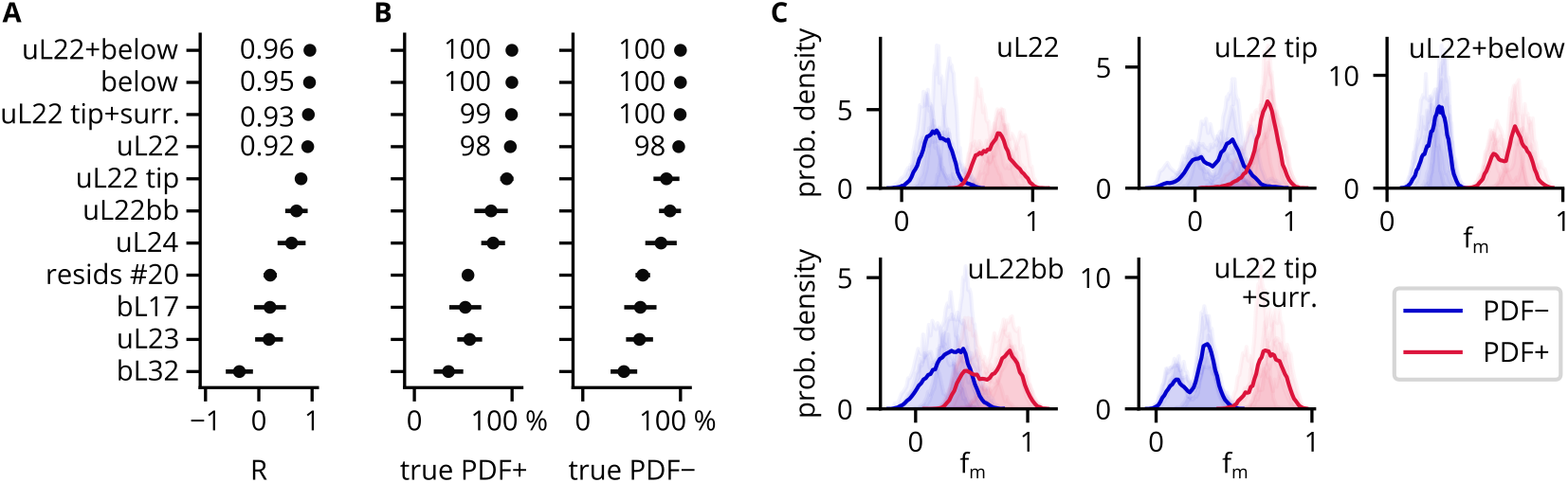
Principal component regression of several ribosomal parts. A) Pearson correlation coefficients *R* obtained as mean values over six validation sets. Errorbars correspond to standard deviations. B) Percentage of correctly classified frames represented by mean values over six validation datasets. Errorbars correspond to standard deviations. C) Probability density functions of *f_m_* calculated for the PDF+ and PDF– components of the validation sets. Light areas represent individual validation sets and the lines stand for aggregated *f_m_* probability densities of all validation sets.

Fig. 3C shows the probability densities of *f_m_* for validation sets. A good model should have the histograms well separated, meaning that PDF+ and PDF– cases are correctly classified by the model. Here, uL22 yielded reasonable predictions. The PCR of uL22 backbone (uL22bb) yielded densities that overlapped more than densities calculated for uL22 including side chains. The model based on uL22bb correctly classified only 78% of PDF+ frames of the validation set. This implies that the uL22 side chains are important for a successful prediction of the ribosome surface status.

To check the trivial case that the good prediction is just due to different conformations of the PDF binding site on the ribosome surface, we tested atoms’ selections excluding the binding site. For instance, the predictions calculated from the uL22 tip alone showed a good correlation (*R* = 0.79 ± 0.08), but with overlapping probability densities. In the validation set, 94% and 85% of frames were classified correctly for the PDF+ and PDF– cases, respectively. Including the nucleotides around the tip improved the model considerably both in terms of the classification (99% of frames correctly classified) as well as correlation (R = 0.93 ± 0.02).

The best model was obtained from a selection of uL22 with a portion of rRNA below the PDF C-terminus included (R = 0.96 ± 0.02) which included the PDF binding site. This selection is denoted *uL22+below.* This model was investigated further to clarify presumed allostery.

### 2.4 Maximally Correlated Motion Explains the Allostery

In PCR, a vector 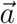 is defined as a linear combination of input PCA eigenvectors 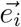

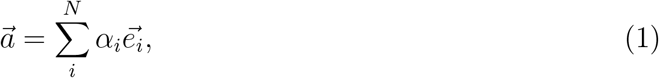

which represents a conformational change best correlated with *f*. The vector 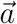 is, in the spirit of FMA,^25^ denoted *maximally correlated motion* (MCM). Thus, the vector components were projected onto the structure revealing sites important for good prediction of PDF presence.

Figure 4A shows the selection uL22+below. The atoms with the highest MCM components are highlighted (see Methods). The results for individual trajectories are shown in Fig. S2.

**Figure 4:**
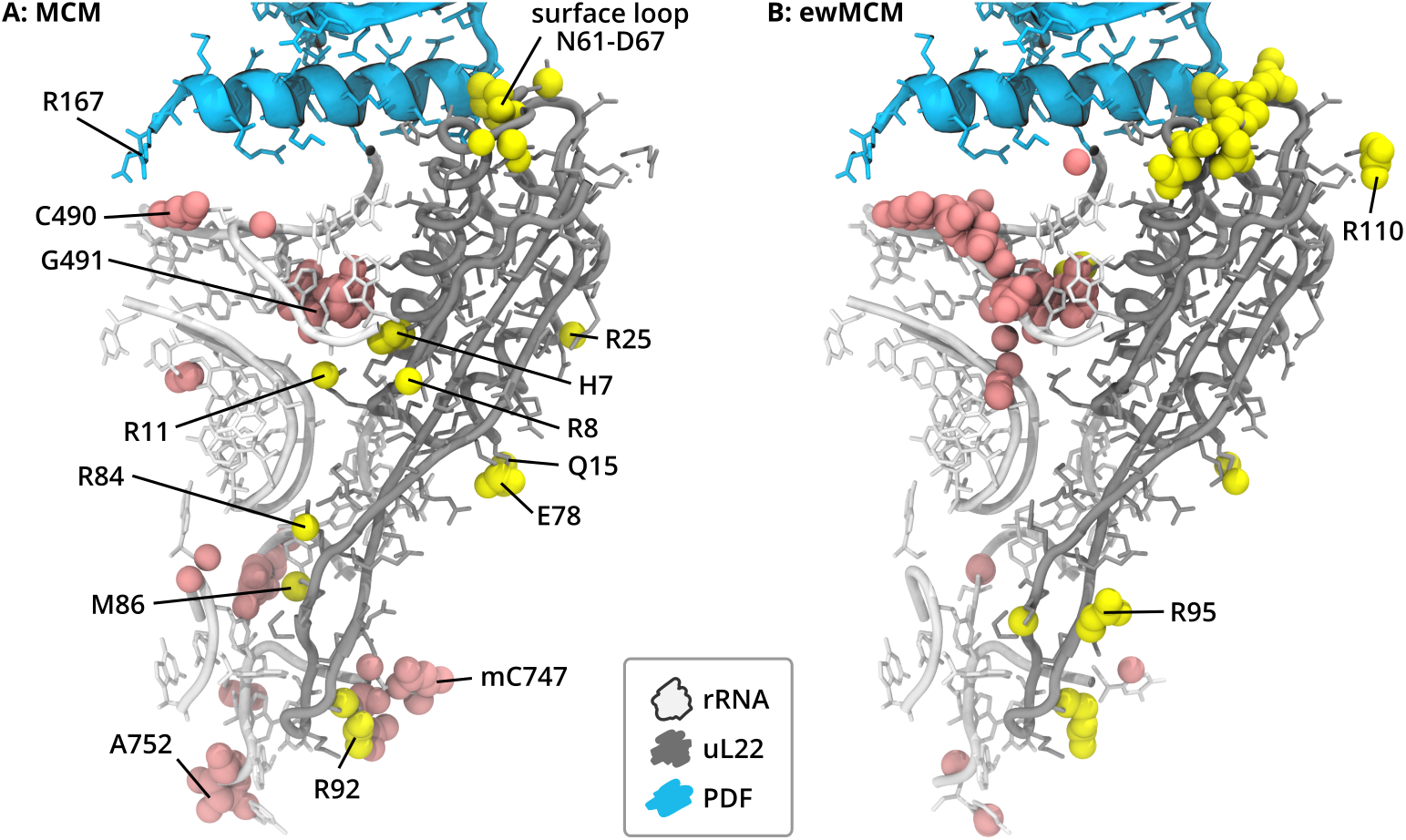
PCR results projected onto the ribosome structure. PDF is shown in cyan, uL22 in gray, a portion of 23S rRNA in white. A consensus selection of atoms contributing the most to the maximally correlated motion (MCM, A), or ensemble-weighted MCM (ewMCM, B) of the *uL22+below* selection are shown as pink and yellow spheres for rRNA and uL22, respectively.

First, the presence or absence of the PDF is connected with a motion of the uL22 surface loop near N61, as indicated by numerous atoms contributing substantially to the MCM. This observation is consistent with the rwRMSD of both experiment- and simulation-based structural models (Fig. 2). Second, the PDF binding is correlated with a motion of a 23S rRNA loop of residues C487 to G493 with the most affected nucleotide being C490. In the bare ribosome, C490 frames the tunnel exit on the side opposite to the tip of uL24. When PDF binds, its C-terminal R167 directly affects the conformation of C490, which likely propagates downwards through the tunnel walls. Apart from the walls, the signal propagates via the interface between uL22 and 23S rRNA, where positive side chains of R8, R11, R25, R84, and neutral H7, Q15 and M86 possess a high MCM component. Finally, the PDF presence and absence is correlated with the motion near the tunnel constriction, namely by the R92 of uL22 and mC747 and A752 of rRNA. It seems likely that the conformational motion of the R11 side chain facilitates the communication between the rRNA loop near the tunnel exit, and the tunnel constriction. Other residues on the boundary of the globular and loop-extension domains of uL22 (R8, Q15, E78) may also contribute.

The Gibbs energy landscape governs the conformational motion of a system of interest and may restrict the MCM determined by PCR. Thus, the MCM may cover motions, which are unavailable for the system at the given temperature. To better understand allostery represented by the MCM, we calculated ensemble-weighted MCM (ewMCM),^25^ which represents “the most probable collective motion that accomplishes a specific displacement along the MCM”.^25^ Therefore here, the ewMCM vector shows the motion most correlated with the presence or absence of PDF *and* observed in the MD simulations.

Figure 4B reveals that the ewMCM overlaps with the MCM in several regions, most notably in the surface loop of uL22, and the rRNA loop below the PDF C-terminal helix. The residues in the exit tunnel like R92 of uL22 or A752 of rRNA contribute too, but the amount of atoms picked by consensus (see Methods) with a high ewMCM contribution is lower in this region than for the MCM. Interestingly, the residues on the boundary between the uL22 globular and loop-extension domains (R11, Q15, etc. in Fig. 4) are missing in ewMCM. It means that on one hand in MD simulations, the motion of H7, R11, or E78 possesses a low variance, so it does not appear in the ewMCM. On the other hand, due to including low-variance PCs for the PCR, the motion of these residues appears in the MCM. Consequently, the allostery triggered by PDF, as identified by PCR, does not fully occur within the time scale of the simulations and remains latent in certain ribosome parts.

## 3 Discussion

We used atomistic MD simulations and a supervised-learning statistical model to understand the effect of PDF binding to the ribosome surface. We based our simulations on several experimental structures. The cryo-EM structure of the ribosome-PDF complex was only partial, because the PDF binding mode was represented by the PDF’s C-terminal *α*-helix.^21^ Our simulations confirmed the binding mode for the full PDF. Unlike cryo-EM, our simulations revealed that due to the extensive conformational flexibility of PDF, the uL22 surface loop may occasionally interact not only with the C-terminal helix, but also with the PDF’s main catalytic domain. Therefore, the structural effect of PDF on the uL22 surface loop was larger in simulations than in experiments. On the other hand, the simulation models were build from the bare ribosome, so slow conformational changes could not occur during the microsecond simulation time. This was the case for several uL22 side chains, most importantly the side chain of R11.

Recently, the kinetic mechanism of the formylmethionine deformylation was described in detail.^26^ The rate limiting step of the deformylation is the dissociation of the deformylated NC from the PDF. The PDF binding to the ribosome is a rapid process. According to the study,^26^ the rate constant is about 40 s^−1^, which is much higher than the overall rate of translation. Our simulations showed a fast and extensive conformational motion of the PDF with respect to the ribosome on the sub-microsecond time scale. It means that, within a single PDF-binding event, the PDF may scan the formylated (unprocessed) NC, which may facilitate the NC recruitment to the PDF catalytic cavity.

The extensive conformational motion of PDF may also contribute to experimentally observed interference of PDF with other protein-biogenesis factors like trigger factor (TF) and methionine aminopeptidase (MAP). A low-resolution cryo-EM structure suggested^22^ that PDF and MAP share a common binding site. When PDF is bound, however, the MAP is able to bind on an alternative site so the N-terminal excision may be accomplished faster. Bornemann et al. argued that the PDF changes the mode of TF binding^27^ on translating ribosomes. The cryo-EM structure of the ribosome-PDF-TF ternary complex confirmed a simultaneous binding of the two factors.^22^ The motion of PDF bound to the ribosome suggests that the PDF may also influence more distant locations on the ribosome surface other than the PDF binding site only.

A major finding of this study was the identification of allostery triggered by the PDF binding to the ribosome. Our simulations and statistical modeling revealed a path propagated from the PDF through the tunnel walls deeper into the tunnel. The cross-validation pointed out that MCM vectors differed: a typical scalar product of MCM vectors from two different validation sets was about 0.6 (Fig. S3). Consequently, the selections of atoms contributing the most to the MCM were different for various validation sets (Fig. S2). Still, our conclusions are independent on the training/validation set: i) uL22 side chains, not the backbone, are correlated with the ribosome surface status, ii) the tunnel constriction including uL22 aminoacid residues and 23S rRNA is correlated with the ribosome surface status, and iii) rRNA surrounding the 23S residue C490 transfers the allosteric signal deeper to the tunnel. The MCM sites overlap with allosteric regions identified by an analysis of multiple sequence alignment of 23S RNAs.^28^ This is true especially about 23S residues between mC747 and A752 located next to the uL22 tip, which were shown to be involved in a long-distance coupling.^28^

The equilibrium constant of PDF-ribosome association is independent – within the experimental uncertainty – of the translation status of the ribosome.^26,29^ Hence, PDF binds equally strongly whether the nascent chain is present in the tunnel or not. This suggests that the allosteric signal is likely unidirectional, from the surface to the interior.

We showed that PDF modulates the structure and dynamics of distant parts of the empty exit tunnel, most notably the tunnel constriction. The constriction contains multiple side chains with a positive charge, some of which were identified as coupled to the PDF binding (R92 of uL22, for instance). Previously, it has been shown that the electrostatics of the exit tunnel affect the rate of NC translocation.^30^ Also, the tunnel near the constriction encourages nascent chain compaction.^17^ The delicate equilibrium of the constriction may affect how nascent peptides translocate through the tunnel. We speculated that this may be especially important early during translation, when the N-terminus of NC diffuses through the tunnel towards the exit. The pulling force later generated by the co-translational folding^31^ is likely missing at this stage, so even small variances of the tunnel walls may support the translocation. PDF may thus ease, though the allosteric modulation of tunnel walls, the N-terminus ejection.

## 4 Materials and Methods

### 4.1 Simulated Systems

The initial structural model of the ribosome–PDF complex was derived from the cryo-EM structure of the *E. coli* ribosome (5AFI,^24^ average resolution of 2.9 Å). The PDF C-terminal helix was added from the available ribosome–helix complex (4V5B,^21^ average resolution of 3.7 Å) by superimposing the large ribosomal subunits. Finally, the PDF was completed using an X-ray structure of an *E. coli* PDF–ligand complex (2AI8,^32^ average resolution of 1.7 Å). The ligand was removed and the C-terminal helices were superimposed. Apart from the ribosome–PDF complex (PDF+), we also simulated the bare ribosome (PDF–).

Each model was placed in a rhombic dodecahedron periodic box of water ensuring a minimal distance of 0.9 nm between the solute and box faces. Structural Mg^2+^ ions were used as found in the cryo-EM ribosome model (5AFI) and the catalytic Ni^2+^ ion was used as in the X-ray (2AI8). The box was neutralized by randomly replacing a sufficient number of water molecules by K^+^ ions. To mimic physiological conditions, excess amounts of MgCl_2_ and KCl were added in order to reach concentrations of 10 and 100 mmol dm^−3^, respectively.

We used the Amber12SB force field^33^ for proteins and canonical nucleotides, Aduri et al. parameters for non-canonical nucleotides,^34^ SPC/E water parameters,^35^ and Joung and Cheatham ion parameters.^36^ The same setup has proven reliable in previous simulation studies of the ribosome.^6,12,37^ The catalytic Ni^2+^ ion was kept in the X-ray conformation using harmonic distance restraints to the four nearest interacting residues (Gln50, Cys90, His132, and His136) using a force constant of 10,000 kJ mol^−1^ nm^−2^.

### 4.2 Molecular Dynamics Simulations

Each system was equilibrated in a series of short simulations. First, the energy of the solvent was minimized in 50,000 steps using the steepest descent algorithm keeping all solute atoms restrained by a harmonic potential with a force constant of 1000 kJ mol^−1^ nm^−2^. Second, the solvent was heated over the course of a 500 ps MD simulation to 300 K using the v-rescale thermostat,^38^ while the solute was kept at 10 K and position restrained. The initial velocities were selected randomly from the Maxwell-Boltzmann distribution at 10 K. The pressure was equilibrated to 1 bar in a 2-ns long MD simulation using the Berendsen barostat.^39^ The position restraints were gradually released and the solute was heated to 300 K during 10 ns simulation. Finally, the production simulations were carried out at a constant temperature of 300 K and a pressure of 1 bar using the v-rescale thermostat and the Parrinello-Rahman barostat.^40^ For each system we generated 4 independent trajectories, 1040 ns each, differing in the initial velocities. For the analyses, the trajectory frames were saved every 200 ps.

In all simulations, the leap-frog algorithm was used to integrate Newton’s equations of motion. All bond lengths were constrained by P-LINCS^41^ and the hydrogen atoms were treated as virtual sites.^42^ This allowed for a time step of 4 fs for the production runs. The long-range electrostatic interactions were treated by the particle-mesh Ewald algorithm^43^ with the direct-space cut-off of 1.0 nm, interpolation order of 4 and 0.12 nm grid spacing. Lennard-Jones interactions were truncated beyond 1.0 nm. All simulations were carried out in the GROMACS 2019 simulation package.^44^

### 4.3 Analyses

Principle component analysis (PCA) of the PDF Cartesian coordinates^45^ was done after superimposing the independent trajectories using C*_α_* and phosphorus atoms of the large subunit with respect to the experimental structure of the bare ribosome (5AFI^24^). From the concatenated trajectories, frames of every 2 ns were used to construct the covariance matrix.

A subset of the eigenvectors of the covariance matrix, i.e. principle components (PCs), of various atoms selections were used for principle component regression (PCR). PCR was previously proposed in the context of biomolecular simulations as functional mode analysis (FMA).^25^ A functional quantity *f* was defined to reflect the presence (*f* = 1) or absence (*f* = 0) of PDF and a statistical model was built to predict f, the predicted values are denoted *f_m_.*

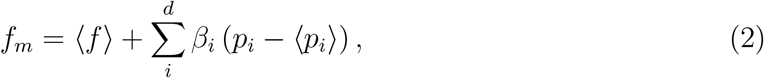

where 〈·〉 denotes an ensemble average, *β_i_* are coefficients determined by the training process, *p_i_* is a projection of the trajectory onto 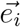 (the PC vector), and the sum runs over the *d* selected PCs.

The model was trained by minimizing the mean-square error (MSE)

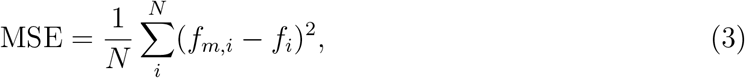

where the sum runs over the *N* frames of the training set. In practice, the problem is solved by maximizing the Pearson correlation coefficient *R*_train_ (justified in Ref. 25) using the training set.

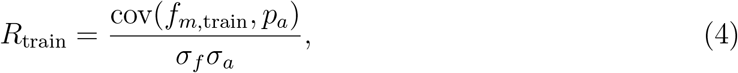

where *σ_f_* and *σ_a_* are standard deviations of *f*_*m*,train_ and *p_a_* (the projection of the trajectories onto 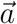, see Eq. 1), respectively, and cov is the covariance.

The model was trained using two PDF+ and two PDF– trajectories, the validation set consisted of the remaining pairs of trajectories. The last 500 ns of each trajectory were used to allow for system relaxation. The model was built using the highest-eigenvalue PCs covering 95% of the total variance, thus the number of dimensions was vastly reduced. The PCA space was constructed only from the training data. Since it was not *a priori* known, which pairs of trajectories should be used for training and validation, we performed crossvalidation. All six possible combinations for training and validation sets available for the four PDF+ and four PDF– trajectories were examined. Then, the quality of the atom selections was assessed using *R*_test_ averaged over the six validation sets. The histograms were calculated for each validation set separately as well as for data aggregated from all validation sets. The MCM (Eq. 1) was calculated for each validation set and a consensus view was obtained as an intersection of the 20% of atoms with the highest MCM contribution.

The ensemble-weighted MCM (ewMCM) vector 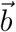 was defined as

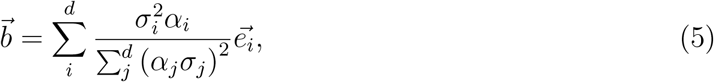

where 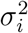 is the variance of the i-th PC and the sums run over *d* selected PCs. ewMCM represents the most probable motion along 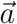 as observed in simulations.^25^

The analyses were done with custom python scripts using the MDAnalysis library.^46,47^ The scripts together with exemplary input and output files are available at https://github.com/mhkoscience/mcgrathh-pdfribo.

## 5 Authors’ contribution

HM performed MD simulations and analyses, MČ built the structural models and helped with the analyses, MHK performed MD simulations and analyses, and with the help of all other authors wrote the manuscript.

## 6 Acknowledgement

We thank Lars V. Bock for invaluable discussions and commenting the manuscript, and Lucie Kubíčková for her help with the simulation system preparation. The research was supported by the Czech Science Foundation (project *19-06479Y*). MČ acknowledges the support of the Internal Grant Agency of UCT Prague (project *403-88-2091*). We thank Max Planck Computing and Data Facility for the computing time.

## Supplementary information

### Input Data and Analysis Scripts

The data necessary to run molecular dynamics simulations, analyses scripts, and exemplary input and output files are available online on https://github.com/mhkoscience/mcgrath-pdfribo.

### Supplementary Figures

**Figure S1:**
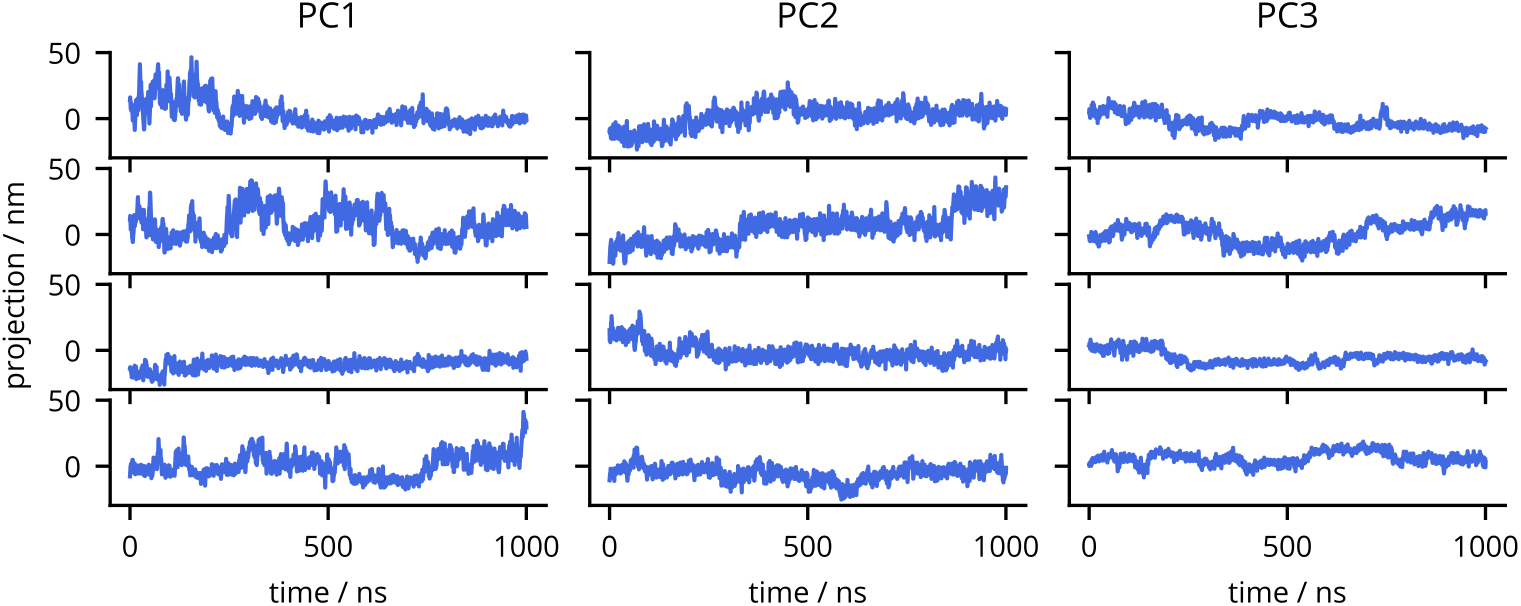
Time evolution of trajectory projections on the first three principle components (PCs).

**Figure S2:**
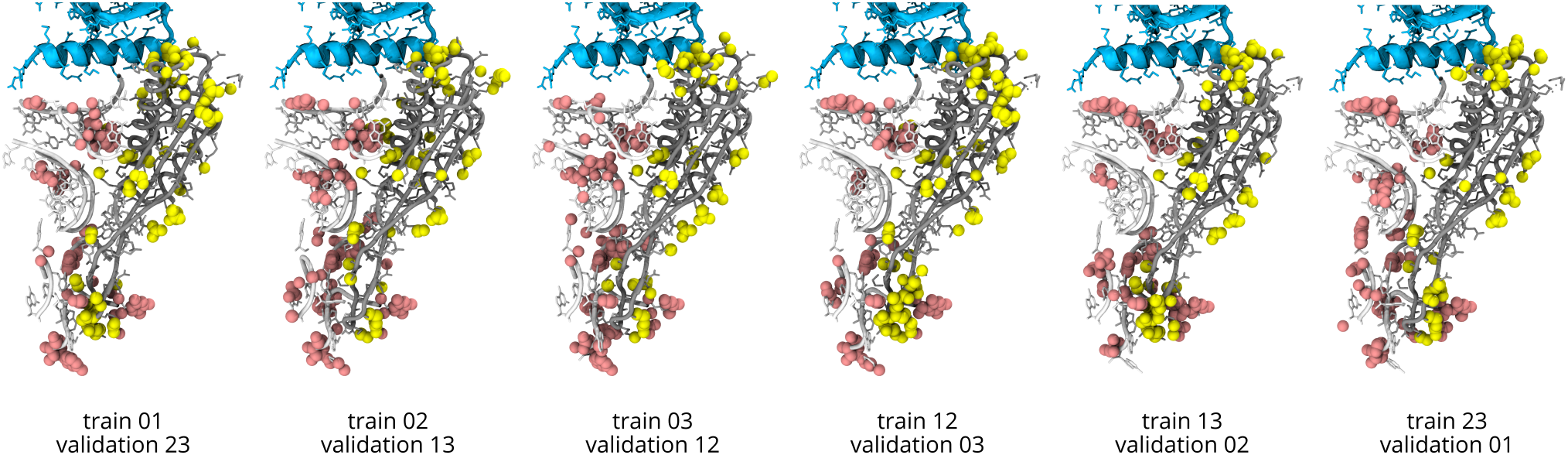
Maximally correlated motion calculated by principle component regression using various training and validation sets. Peptide deformylase is shown in cyan, ribosomal protein uL22 in gray, a portion of 23S rRNA in white. The 20% of atoms with the highest MCM contribution of the validation set is shown in pink for rRNA and yellow for the uL22.

**Figure S3:**
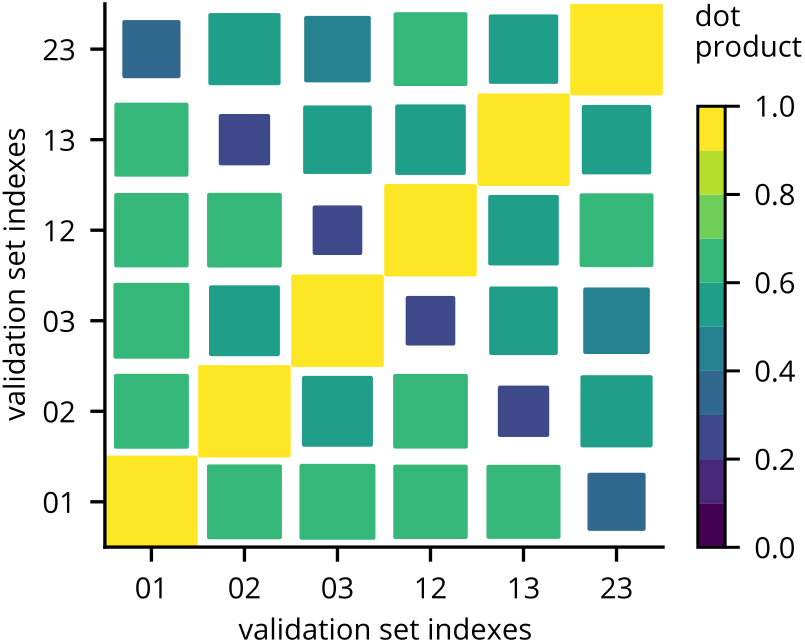
Dot products of MCM vectors calculated for various validation sets. The area of the squares is proportional to the dot product.

